# Hemodynamic traveling waves transport musical codes faster than language codes via shared bilateral streams

**DOI:** 10.1101/2025.03.04.641396

**Authors:** Ruey-Song Huang, Ci-Jun Gao, Ut Meng Lei, Teng Ieng Leong, Jian Hwee Ang, Cheok Teng Leong, Chi Un Choi, Martin I. Sereno, Defeng Li, Victoria Lai Cheng Lei

## Abstract

What takes precedence in our brain: music or language? Here we used rapid phase-encoded fMRI to explore spatiotemporal brain dynamics during naturalistic music and language tasks, seeking to track down the entire perception-to-production process and compare the speed of information transfer between the two domains. We found largely shared streams of traveling waves bilaterally, along which musical codes were transported faster than language codes for identical visual input.

## Main text

Music and language, two universal human attributes, have captivated scholars from ancient times to modern neuroscience^1–6^. Functional magnetic resonance imaging (fMRI) studies have revealed intriguing neural overlaps and differences between music and language processing^6–21^. However, the temporal dynamics of information flows in both domains remain largely unexplored^22^. Several challenges persist in comparing real-time music and language processing using fMRI. First, contrast-based fMRI designs and analyses cannot reveal the timing and directions of information flows across the brain. Second, matching stimulus properties between music and language is complex, with discrepancies in stimuli and tasks inevitably leading to differences in brain activations^9,23^. Third, head motion artifacts and scanner noise pose significant challenges in fMRI experiments involving overt production. Designs are usually limited to passive perception of stimuli^13,20^, covert production^24,25^, or overt production with sparse fMRI sampling^16^.

In this study, we used rapid phase-encoded fMRI^26,27^ to capture the dynamic flows of musical codes and language codes via hemodynamic traveling waves across the cortical surface during naturalistic perception and overt production tasks. For tasks involving reading, Western Arabic numerals were presented and interpreted either as digits (basic language units), or as numbered musical notation (basic music units). Subjects were scanned continuously while reading the digits or musical notes silently, and then reciting the digits (in Mandarin), singing the musical notes, or playing the keyboard (with the right hand), with real-time auditory feedback through headphones (Fig. 1; Methods). In the digit reading-reciting task, subjects read and memorized seven digits from 0 to 4 s, recited them from 4 to 8 s, and rested from 8 to 16 s in each cycle (Fig. 1c), which repeated 16 times in each scan. The amplitude (signal-to-noise ratio) and phase of signals at 16 cycles per scan in each voxel were color-coded and rendered on individual cortical surfaces^26–29^ (Fig. 1d; Methods). Time courses of periodic activations with different delays were averaged within each selected surface-based regions of interest (sROIs; Fig. 1e,f). Surge profiles^27^ reveal when the hemodynamic traveling waves rise, peak, and subside in these sROIs (Fig. 1g; Methods).

**Fig. 1.**
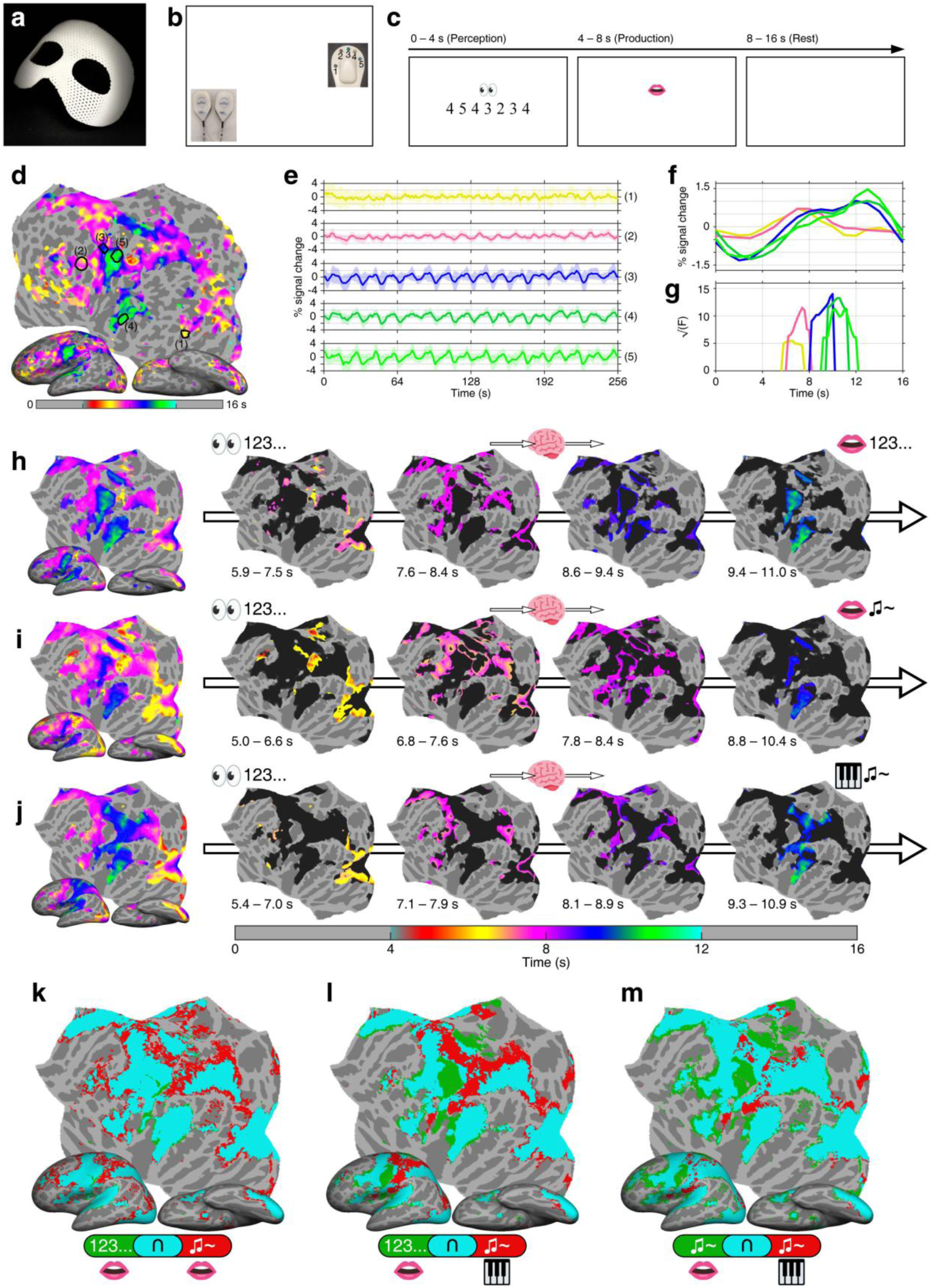
**a**, A custom-molded mask for preventing head motion. **b,** Experimental setup for language and music tasks. The participant wore a mask and noise-cancellation headphones (left inset), with a microphone above the mouth. A 5-digit keypad (right inset) was placed under the right hand. **c,** Timeline of a digit reading-reciting task. **d,** Phase-encoded activations (F_(2,230)_ = 25.492, *P* < 10^-10^, uncorrected) in the left hemisphere of a representative subject, where the colorbar indicates different activation phases during the task. **e,** Average and standard deviation of voxel time courses in five selected sROIs. **f,** Average time courses within a 16-s period in five sROIs. **g,** Surge profiles in five sROIs. **h**, **i**, and **j**, Group-average maps of phase-encoded activations (*n* = 21, *F*_(2,40)_ = 5.18, *P* < 0.01, cluster corrected) in the left hemisphere for the digit reading-reciting task, note reading-singing task, and note reading-playing task, respectively. **k**, **l**, and **m**, Conjunction maps illustrating overlaps between tasks (see percentages of overlaps in Supplementary Table 1). Green or red: regions activated by a single task; Cyan: regions activated by both tasks.

Fig. 1h-j show group-average phase-encoded activations in the left hemisphere for three tasks involving reading. Each map is divided into four phases of activations (see animated traveling waves with continuous phases in Supplementary Videos 1, 2, and 3; Methods). Initially, the perception and encoding of input activated both the ventral and dorsal visual streams^27–29^ (second panels, reddish and yellowish regions). Subsequently, the assembly and storage of codes activated regions in the intraparietal sulcus (IPS), superior parietal lobule (SPL), frontal operculum and anterior ventral insular (FOP/AVI), dorsolateral prefrontal cortex (dlPFC), dorsal premotor cortex (PMd), and dorsomedial frontal auditory field (dmFAF^27,28^) (third panels, pinkish regions). Next, motor planning activated regions in the inferior and superior parietal lobule (IPL/SPL), supplementary motor area (SMA), pre-SMA, rostral middle frontal gyrus (rMFG), ventral premotor cortex (PMv), and posterior superior temporal gyrus (STG) (fourth panels, purplish and bluish regions). Lastly, overt production (reciting or singing) with self-monitoring engaged the articulatory and respiratory areas in primary sensorimotor cortex (MI/SI), Sylvian parietal temporal area (Spt^25,30^), and auditory cortex (fifth panels, bluish and greenish regions). Along with activations in Spt and auditory cortex, playing notes activated manual control regions^28,29^, including hand representations in MI/SI, anterior intraparietal area (AIP), SPL, and secondary somatosensory cortex (SII).

The comparison between the reading-reciting and reading-singing maps shows significant overlaps in the visual, auditory, posterior parietal, sensorimotor, and frontal cortices (Fig. 1k and Supplementary Table 1; Methods). Interestingly, the reading-singing task also activated manual control regions, including MI/SI, AIP, and SPL^28,29^, during the perception and encoding of musical notes (Fig. 1i, second panel). The reading-reciting and reading-playing maps show significant overlaps but differ in articulatory and respiratory areas (greenish) associated with reciting and hand movement areas (reddish) associated with playing (Fig. 1l). The comparison between the reading-singing and reading-playing maps (Fig. 1m) is largely similar to that in Fig. 1l, except in manual control regions.

Fig. 2a-c show group-average phase-encoded activations in the left hemisphere for three tasks involving listening to spoken digits or musical notes (Supplementary Videos 4, 5, and 6; Methods). Initially, the perception and encoding of input activated the primary auditory cortex (A1), anterior STG, superior temporal sulcus (STS), parietal ventral and secondary somatosensory area (PV/S2)^28,29^, 45aud^28^, and dlPFC^28^ (second panels, reddish and yellowish regions). Subsequently, the transformation and storage of language or music codes activated the association auditory cortex, polysensory zone (PZ)^28,29^, dmFAF^28^, pre-SMA, SMA, posterior dlPFC, FOP/AVI, PMd, and IPS (third panels, pinkish regions). Next, motor planning activated frontal and parietal operculum, IPL, Spt, PMv, rMFG, and SMA (fourth panels, purplish and bluish regions). Lastly, reciting and singing activated rMFG and articulatory and respiratory areas in MI/SI, while playing activated manual control regions (fifth panels, bluish and greenish regions). Spt was activated during production and self-monitoring in all three tasks. Significant cortical overlaps are evident in the conjunction map comparing the reciting and singing tasks (Fig. 2d; Supplementary Table 1), with the singing activated more in PMd. Distinctions between the reciting and playing tasks are identifiable in regions involved in vocalization and hand movements (Fig. 2e). The comparison between singing and playing tasks exhibited a similar pattern, with greater overlaps in PMd (Fig. 2f).

**Fig. 2.**
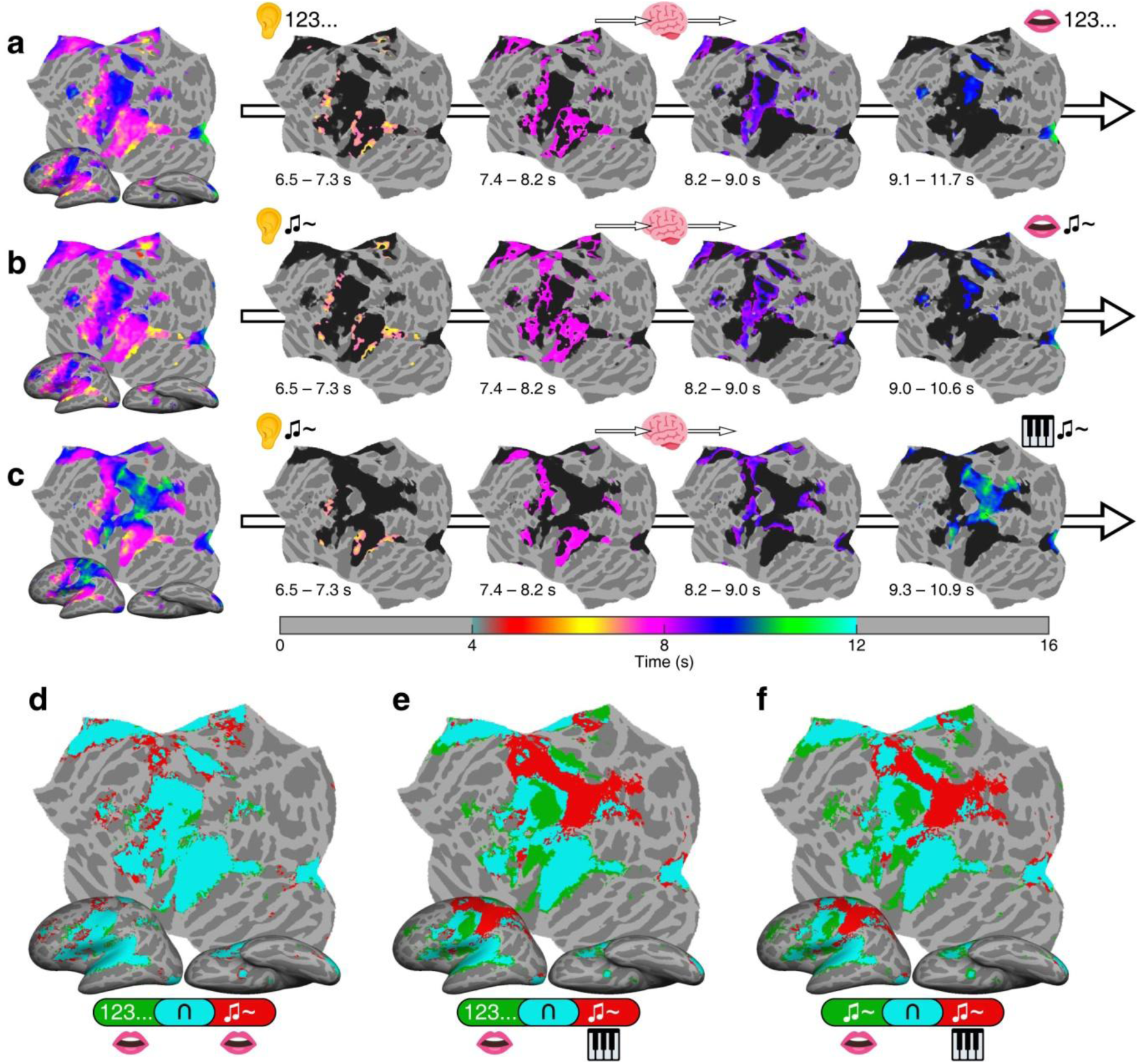
**a**, **b**, and **c**, Group-average maps of phase-encoded activations (*n* = 21, *F*_(2,40)_ = 5.18, *P* < 0.01, cluster corrected) in the left hemisphere for the digit listening-reciting task, note listening-singing task, and note listening-playing task. **d**, **e**, and **f**, Conjunction maps illustrating overlaps between tasks (Supplementary Table 1). Green or red: regions activated by a single task; Cyan: regions activated by both tasks.

The overall spatiotemporal patterns in phase-encoded activation maps are bilaterally symmetric across all reading and listening tasks, except that the right hemisphere does not show activations in hand representations in MI/SI during playing tasks involving only the right hand (Extended Data Figs. 1 and 2). Furthermore, the conjunction maps reveal that PMd and AIP in the right hemisphere were involved in the listening-playing task but not in the reciting and singing tasks (Extended Data Fig. 2e,f).

The surge profiles in Fig. 3a (upper panels; Methods) show the overall distribution of activation phases in the left hemisphere for three tasks involving reading. Whole-hemispheric activations in the singing task rose more rapidly than the reciting task, while activations in the playing task rose slightly slower than the singing task but faster than the reciting task.

**Fig. 3.**
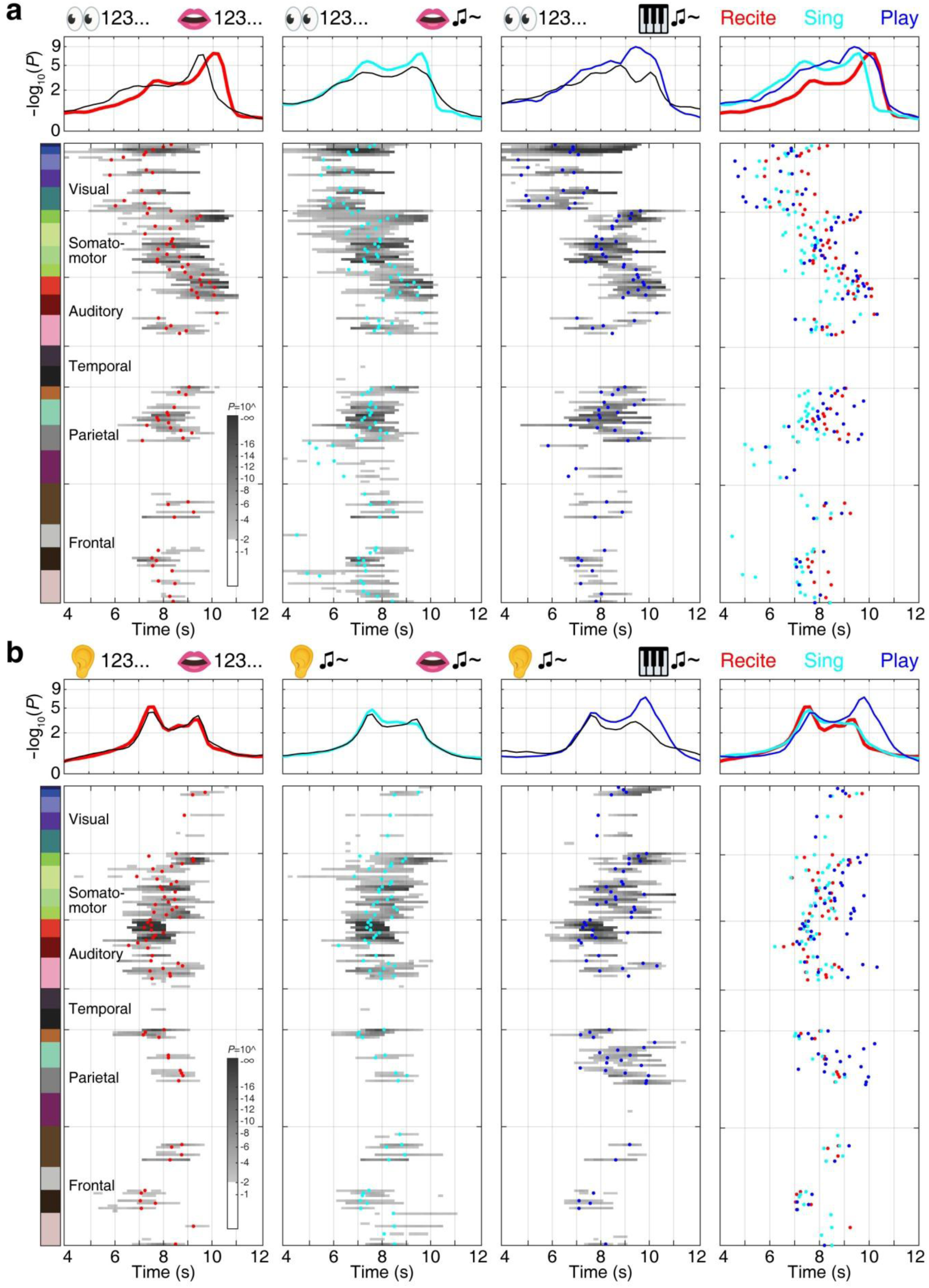
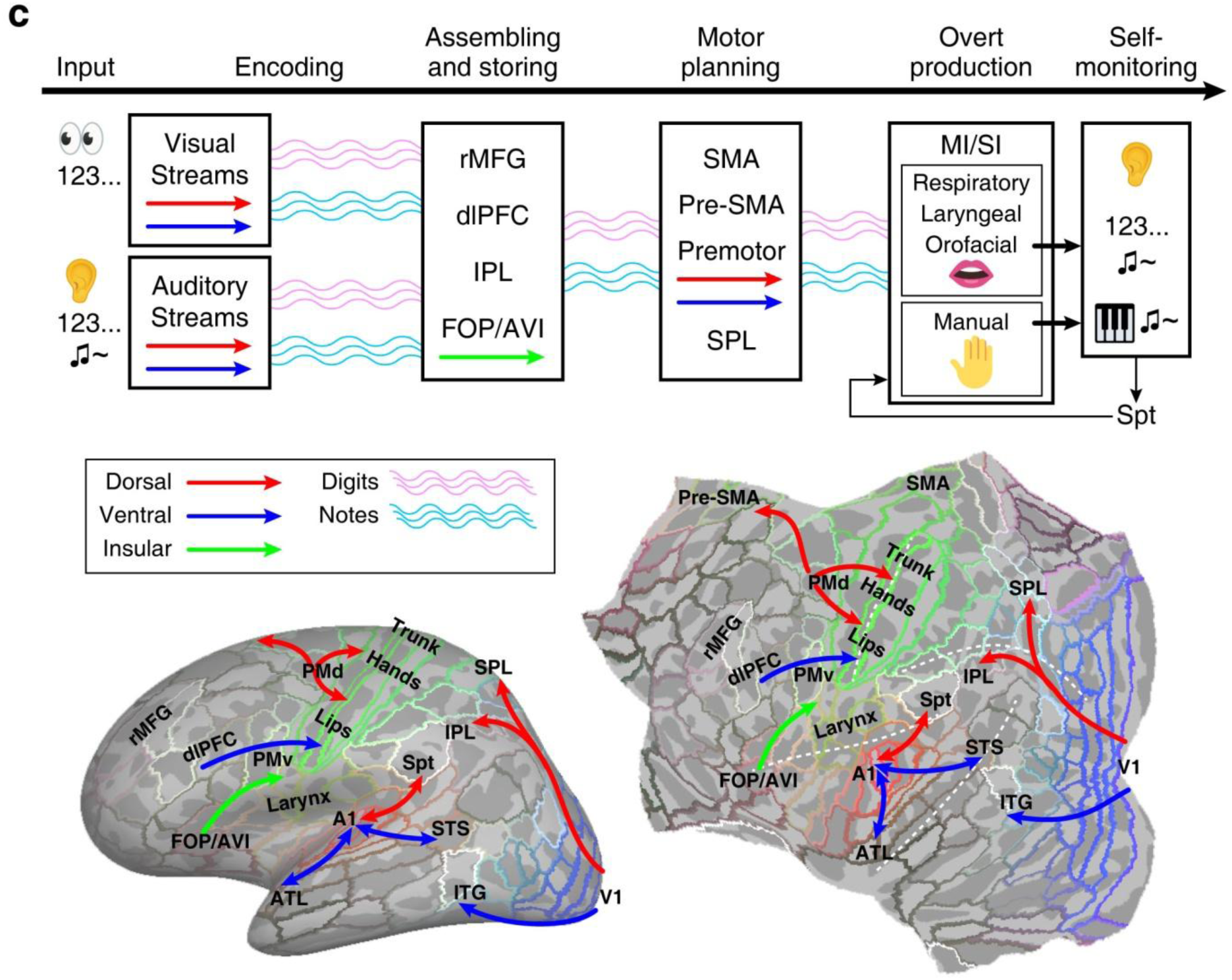
**a**, Surge profiles (upper panels) and Gantt charts (lower panels) illustrate activations in the left hemisphere during the reading-reciting (red), reading-singing (cyan), and reading-playing (blue) tasks. Each black curve displays the surge profile of overall activations in the right hemisphere for each task. The surge height at each time point is computed by -log10(*P*-value), e.g., a surge height of 2 indicates *P* = 0.01 (see Methods). Each grayscale bar in the Gantt chart indicates the surge profile of an sROI, highlighting above-threshold portions (surge height > 2; *F*_(2,230)_ = 4.7, *P* < 0.01). The colorbar indicates 22 groups of sROIs listed in Supplementary Table 3. The dot on each bar indicates the mean phase of vertices within each sROI. The rightmost panels provide comparisons of surge profiles and sROI mean phases across three tasks. **b,** Surge profiles (upper panels) and Gantt charts (lower panels) illustrate activations in the left hemisphere during the listening-reciting (red), listening-singing (cyan), and listening-playing (blue) tasks. All conventions follow those of Fig. 3a. **c**, (Top panel) A neural logistics model for language and music processing, outlining a sequence from perception to production. Wavy lines indicate that language codes (digits) and musical codes (notes) are transported by traveling waves from one process to the next. (Lower panel) Streams of traveling waves in visual, auditory, premotor, and insular cortices, depicted on inflated and flattened surfaces of the left hemisphere.

Furthermore, activations in the singing task subsided faster than the reciting or playing tasks during the production phase. Compared with the left hemisphere, activations in the reciting task rose and subsided faster with comparable amplitudes in the right hemisphere (black curves in the upper-left panels of Fig. 3a). In the singing and playing tasks, the surge profiles were comparable between the hemispheres, with the right hemisphere exhibiting lower amplitudes (Extended Data Fig. 3a).

The surge profiles in Fig. 3b show that left-hemisphere activations rose approximately at the same time across all tasks involving listening. Activation amplitudes were comparable bilaterally in the reciting and singing tasks. However, the left-hemisphere surge profile in the playing task peaked and declined later, related to right-hand movements. In contrast, the right-hemisphere surge profiles did not show activations in the hand representations in MI/SI during the production phase (Extended Data Fig. 3b). Differences in surge profiles were noticeable in the earlier phases of the tasks involving reading (Fig. 3a). However, the surge profiles were comparable between tasks involving listening, except in the production phase of the listening-playing task (Fig. 3b).

The Gantt charts in Fig. 3a,b (lower panels) compare the surge profile (grey-dark bar) and mean activation phase (*θ_sROI_*; dot) in each sROI in the left hemisphere across tasks (see sROI maps based on HCP-MMP1.0 parcellation in Extended Data Figs. 4 and 5; Methods). Across sROIs, the average of *θ_sROI_* is significantly earlier in the reading-singing task compared with the reading-reciting task (*F*_(1,189)_ = 38.31, *P* = 3.65×10^-9^; Extended Data Fig. 6; Methods). In the listening-singing task, the average of *θ_sROI_* is slightly earlier than the listening-reciting task, but the difference is not statistically significant (*F*_(1,139)_ = 0.55, *P* = 0.46). The average of *θ_sROI_* in the reading-playing task is significantly later than the reading-singing task (*F*_(1,198)_ = 15.75, *P* = 0.0001). Similarly, when comparing with the listening-singing task, the average of *θ_sROI_* in the listening-playing task is significantly later (*F*_(1,140)_ = 20.58, *P* = 0.00001).

Compared with the left hemisphere, the distributions of *θ_sROI_* in the right hemisphere do not show a significant difference between tasks (*P* > 0.01; Extended Data Fig. 6). Furthermore, the left hemisphere exhibits greater dominance in the activation maps of both music and language tasks, as shown by the laterality index (LI) maps (Extended Data Fig. 5 and Supplementary Table 3; Methods).

Both the Gantt charts and traveling wave videos reveal how spatiotemporal hemodynamic activations propagate across regions in the visual, parietal, insular, frontal, somatomotor, and auditory cortices (Fig. 3a,b, Extended Data Fig. 3, and Supplementary Videos 1-6). Here, we propose that the neural logistics model^27,28^ for language processing is also applicable for transporting musical codes through multimodal streams of traveling waves across the brain (Fig. 3c). In the reading-reciting task (Supplementary Video 1), for example, printed language codes (digits) are processed through both the dorsal and ventral visual streams, transformed and assembled into a sequence of phonological codes, and temporarily stored in working memory. These verbal codes are then transformed into motor codes, transported through the frontal opercular-insular cortex, ventral and dorsal premotor cortex^31^, and supplementary motor area, delivered to the respiratory, laryngeal, and orofacial areas in MI/SI^28,29^, and finally received as auditory codes for self-monitoring through Spt. Similarly, in the reading-singing task (Supplementary Video 2), Western Arabic numerals are encoded through both visual streams, transformed and assembled into musical codes (notes), and then processed and transported through the same logistics streams for language codes—but faster. Furthermore, the reading-playing task shares streams with the reciting and singing tasks during the early phases, but motor codes are eventually delivered to manual control regions during the production phase (Supplementary Video 3).

In tasks involving listening, spoken digits or piano tones are received by A1 and transported through the dorsal and ventral auditory streams^27,32,33^, and then assembled and stored directly in their original auditory forms in the working memory (Fig. 3c; Supplementary Videos 4-6). No significant difference in the average of *θ_sROI_* was found between listening-reciting and listening-singing tasks (Fig. 3b; Extended Data Fig. 6). In contrast, in reading-reciting and reading-singing tasks, where the stimuli were identical (both presented as Western Arabic numerals), musical codes (notes) were processed faster than language codes (digits) through shared logistics streams (Fig. 3a, upper panels; Fig. 3c). The discrepancy in processing speed between reading and listening tasks may arise from the different processing load of transforming, transporting, and delivering codes.

In summary, we compared spatiotemporal activation patterns between music and language processing by using precisely paralleled stimuli and tasks and overcoming the technical challenges of overt production in the MRI scanner. While confirming largely shared neural resources between music and language^2,3,12^, rapid phase-encoded fMRI revealed hemodynamic travelling waves that illustrate how neural information flows in time and space via bilateral activation streams. In tasks involving visual input, where the basic language and music prompts were identical in form (Western Arabic numerals), we found that waves of activations during musical tasks travelled faster than those for language tasks along overlapping streams, suggesting that the information processing load for basic music tasks is lighter. The similar flow directions but different processing speeds of neural streams for integrating and transferring multimodal information in these two tasks suggest that music processing is closely related to language processing and may have even served as partial scaffold for it. This supports the speculation that communicative musical abilities may have predated linguistic abilities^34,35^.

## Methods

### Participants

Twenty-one native Mandarin speakers (10 males, 11 females; average age 20.9 ±2.6 years) participated in this study. All participants had over three years of piano training (average starting age 9.3 ±5.1 years), normal or corrected-to-normal vision, and no history of neurological impairment. Written informed consent was obtained from all subjects in accordance with protocols approved by the University of Macau’s research ethics committee.

### Experimental design

Each subject participated in twelve 256-s functional scans in an fMRI session, involving six different tasks using phase-encoded fMRI designs^26–29^, including reading-reciting, reading-singing, reading-playing, listening-reciting, listening-singing, and listening-playing tasks. Each task was repeated across two non-consecutive scans, with each scan comprising sixteen 16-s trials. In the perception phase of each trial (0 to 4 s; Fig. 1c, left panel), the subject silently read seven written digits (randomized between 1 and 5) for reading tasks, or listened to seven spoken digits (randomized between 1 and 5) in Mandarin or seven diatonic piano tones (randomized between 1[C3] and 5[G3]) for listening tasks. Sixteen sequences of spoken digits were generated by Microsoft Azure AI (https://azure.microsoft.com), and 32 sequences of piano tones were generated using FreePiano software (https://freepiano.tiwb.com). Both auditory stimuli were recorded using Audacity software (https://audacityteam.org). In the production phase of each trial (4 to 8 s; middle panel, Fig. 1c), upon being prompted by a visual cue (an icon of a mouth or a mini keyboard), subjects recited (in Mandarin), sang (hummed), or played the memorized stimuli. During the rest phase of each trial (8 to 16 s; right panel, Fig. 1c), subjects viewed a blank screen until the onset of the next trial. Subjects kept their eyes open throughout each functional scan.

### Experimental setup

Before an fMRI session, each subject underwent brief training in an MRI simulator (Shenzhen Sinorad Medical Electronics Co., Ltd.). To prevent head movements during tasks involving vocalization, subjects wore an individualized facial mask molded from thermoplastic sheets (Fig. 1a; 1.6 mm H-board, Sun Medical Products Co., Ltd.). Wearing a mask and a pair of headphones, subjects practiced maintaining head stability while engaging in reciting and singing exercises in the MRI simulator. Real-time monitoring of head movements was facilitated by a motion sensor (MoTrak, Psychology Software Tools, Inc.) affixed to the subject’s forehead, with auditory feedback delivered through the headphones upon exceeding predefined thresholds for translation (1 mm) or rotation (1°).

During the fMRI experimental setup, subjects wore earplugs and MR-compatible noise-cancellation headphones (OptoActive II, OptoAcoustics Ltd.) and lay supine within a head coil filled with deformable resin clay. An MR-compatible microphone (OptoAcoustics Ltd.) was positioned near their mouths for voice recording and real-time auditory feedback via the headphones. A rear-mirror atop the head coil permitted visualization of stimuli on a 40-inch MR-compatible LCD monitor (InroomViewingDevice, NordicNeuroLab AS). An MR-compatible keypad (Fig. 1b; Shenzhen Sinorad Medical Electronics Co., Ltd.) under the subject’s right hand was used to record music playing responses. The buttons under five fingers were mapped to C3 to G3 keys on a virtual piano (FreePiano software), which generated piano tones in real time through the headphones. Visual and auditory stimulus presentation and response recording were managed using Experiment Builder (SR Research Ltd.), awaiting initiation signals (―s‖ key) from the SyncBox (NordicNeuroLab AS) before the commencement of each functional scan. The subjects’ auditory output (speaking, singing, and piano playing) and TTL pulses from the MRI scanner were recorded continuously during each 256-s scan using OptiMRI 3.1 Software (OptoAcoustics Ltd.). The timings of response onset and offset of each trial were identified manually from the soundtracks using Audacity software. The group-average response time, duration, and accuracy are summarized in Supplementary Table 2.

### Image acquisition

Functional and structural brain images were acquired using a 32-channel head coil in a Siemens MAGNETOM Prisma 3T MRI scanner at the Centre for Cognitive and Brain Sciences, University of Macau. Each fMRI session (∼2 hours) consisted of twelve functional scans and two structural scans. Each functional scan was acquired using a blipped-CAIPIRINHA simultaneous multi-slice (SMS), single-shot echo planar imaging (EPI) sequence (acceleration factor: 5; interleaved ascending slices; TR: 1000 ms; TE: 30 ms; flip angle: 60°; 55 axial slices; field of view: 192×192 mm; matrix size: 64×64; voxel size: 3×3×3 mm; bandwidth: 2368 Hz/Px; 256 TR per image; dummy: 6 TR; scan time: 256 s). Two sets of T1-weighted structural images were acquired using an MPRAGE sequence (TR: 2300 ms; TE: 2.26 ms; TI: 900 ms; Flip angle: 8°; 256 axial slices; field of view: 256×256 mm; matrix size: 256×256; voxel size: 1×1×1 mm; bandwidth: 200 Hz/Px; scan time: 234 s) with the same slice center and orientation of the functional images.

### Image preprocessing

Functional images (*.ima files) were converted to the Analysis of Functional NeuroImages (AFNI; https://afni.nimh.nih.gov/) BRIK format using AFNI *to3d* program. All BRIK files were registered with the first volume (target) of the seventh functional scan and corrected for motion using AFNI’s *3dvolreg* program. With head restraining measures, including custom-molded masks and deformable filling inside the head coil, no subject showed major motion artifacts in functional images.

Bilateral cortical surfaces of each subject’s brain were reconstructed from the average of two sets of structural images using FreeSurfer 7.2^36,37^ (https://surfer.nmr.mgh.harvard.edu/). All motion-corrected and slice-timing-corrected functional images were aligned with the structural images acquired right before the seventh functional scan and subsequently registered with each subject’s cortical surfaces using the *csurf* package^28^ (https://pages.ucsd.edu/~msereno/csurf/ or https://mri.sdsu.edu/sereno/csurf/), which includes programs for functional image analyses as detailed below.

### Fourier-based analyses

For each functional dataset (64×64×55 voxels, 256 TR) of each subject, the time series *x_m_*(*t*) of voxel *m* was analyzed with a 256-point discrete Fourier transform^26–28,38–40^:

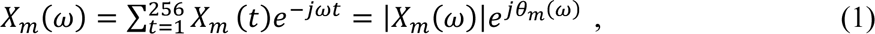

where *X_m_*(*ω*) is the Fourier component at frequency *ω* between 0-127 cycles per scan, and |*X_m_*(*ω*)| and *θ_m_*(*ω*) are its amplitude and phase. The task frequency is defined as *ω_s_* (16 cycles per scan), at which the BOLD signal exhibits periodic fluctuations in response to periodic stimuli and tasks. The remaining non-task frequencies are defined as *ω_n_*. The signal and noise are defined as the Fourier components at frequencies *ω_s_* and *ω_n_*, respectively. The statistical significance of periodic fluctuations of the BOLD signal in voxel *m* is evaluated by a signal-to-noise ratio:

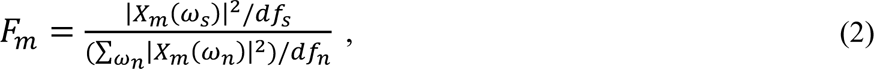

where *df_s_* = 2 and *df_n_* = 230 are the degrees of freedom of signal and noise, respectively. The *P*-value of this *F*-ratio is estimated by the cumulative distribution function *F*_(2, 230)_ = *F*(*F_m_*; *df_s_*, *df_n_*)^26,27,38–40^ . A complex *F*-value, 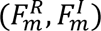, incorporating both the *F*-value and the phase, *θ_m_*(*ω_s_*), of each voxel was obtained by 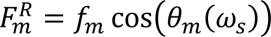 and 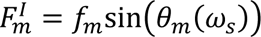, where *f_m_* is the square root of *F_m_*. Voxels containing strong periodic activations at the task frequency (*ω_s_* = 16 cycles per scan, *F*_(2, 230)_ > 4.7, *P* < 0.01, uncorrected) were retained for each functional dataset and displayed on each subject’s cortical surfaces using *csurf*. The phases of these voxels were color-coded between 0.5π and 1.5π, which is equivalent to a range between 4 and 12 s (see colorbar below Fig. 1j).

For each subject *S*, the complex *F*-values in voxel *m* at location (x, y, z) were vector-averaged (voxel-wise) across two scans, k ={1, 2}, of the same task using:

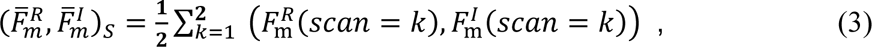

which was carried out by the ―Combine 3D Phase Statistics‖ function in *csurf*. The resulting single-subject average *F*-values, 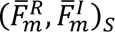, were then projected onto vertex *v* on the cortical surfaces of subject *S*, yielding a map of 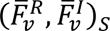.

The spherical averaging method^26,27,38–42^ (“Cross Session Spherical Average‖ function in *csurf*) was used to obtain surface-based group-average maps for each task. First, each single-subject vector-average map, 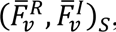, was resampled to a common spherical coordinate system using FreeSurfer’s *mri_surf2surf* program (https://freesurfer.net/fswiki/mri_surf2surf). Second, the complex *F*-values of each vertex *v* on the common spherical surface were vector-averaged (vertex-wise) across all subjects using:

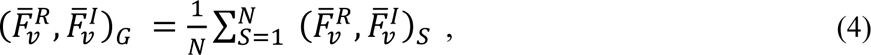

which yielded a map of group-average complex values, 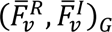, for each task.

The *F*-value of each vertex was obtained by:

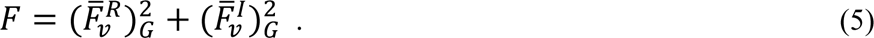

Vertices with significant activations (*F*_(2,230)_ > 4.7, *P* < 0.01) in single-subject surfaces were further tested across subjects (*n* =21, *F*_(2,40)_ > 5.18, *P* < 0.01), and corrected for multiple comparisons using surface-based cluster-size exclusion^40,42^ (cluster = 64 mm^2^, *P* = 0.05, corrected). The phase and amplitude of 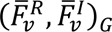 in the cluster-corrected maps were displayed on the inflated and flattened surfaces of FreeSurfer *fsaverage* (see left hemisphere maps in Figs. 1h-j and 2a-c; right hemisphere maps in Extended Data Figs. 1a-c and 2a-c).

### Conjunction maps

A surface-based conjunction map (Fig. 1k-m and Fig. 2d-f; Extended Data Figs. 1d-f and 2d-f) was created for comparing the activation extent between each pair of maps (leftmost panels in Figs. 1h-j and 2a-c and in Extended Data Figs. 1a-c and 2a-c). At the same statistical threshold (*F*_(2,230)_ > 4.7, *P* < 0.01, uncorrected), a vertex on the *fsaverage* surface is colored as follows: (1) green for significant activation only in task 1 (e.g., reading-reciting); (2) red for significant activation only in task 2 (e.g., reading-singing); and (3) cyan for significant activation in both tasks (i.e., overlap). The percentages of vertices activated by a single task or both tasks are summarized in Supplementary Table 1. For example, among the vertices in the left hemisphere activated by the read-reciting task, 95% were also activated by the reading-singing task. However, among the vertices in the left hemisphere activated by the read-singing task, only 65.7% were also activated by the reading-reciting task.

### Surface-based regions of interest

To compare activation patterns between tasks, we subdivided each group-average map (leftmost panels in Fig. 1h-j and Fig. 2a-c, and in Extended Data Fig. 1a-c and Fig. 2a-c) into 180 surface-based regions of interest (sROIs) in each hemisphere according to the HCP-MMP1.0 parcellation^43,44^ (Extended Data Fig. 4; Supplementary Table 3).

For each task, we computed *Q*_LH_ (left hemisphere) or *Q*_RH_ (right hemisphere) as the ratio of the count of vertices with significant activations (*F*_(2,230)_ > 7.1, *P* < 0.001, uncorrected) to the total number of vertices within each sROI. A laterality index^45^ (*LI*) (Extended Data Fig. 5; Supplementary Table 3) was computed for each pair of bilaterally symmetric sROIs by:

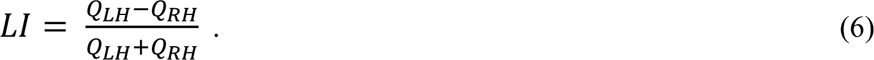

### Surge profiles

A surge profile reveals the timing of arrival, peak, decline, and latency of hemodynamic traveling waves within a brain region during an event or a task^27^. For each task, a surge profile was estimated from the distribution of group-average complex *F*-values, 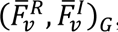, at all vertices within each hemisphere or each sROI (Extended Data Fig. 4) as follows. First, the phase of each vertex *v* was obtained by 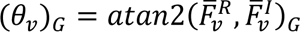 in Matlab software. The complex plane was then divided into 80 equally spaced bins (*d*, time delays) between 0°(0 s) and 360°(16 s). A total of *V* vertices were found with phases, (*θ_v_*)*_G_*, falling within a moving sector centered at bin *d* ={4.5°, 9°, …, 360°}, equivalent to {0.2, 0.4, …, 16.0 s}, where the sector range is 9°(0.4 s) and the step is 4.5°(0.2 s). The vector-average of complex *F-*values of *D* vertices in the moving sector [*d*-4.5°, *d*+4.5°] centered at bin *d* was obtained by:

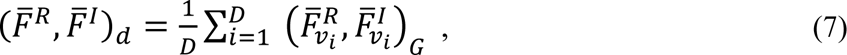

The magnitude of 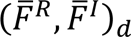 was obtained by:

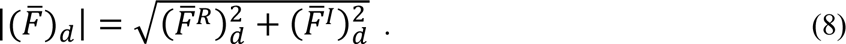

A *P*-value was estimated for each 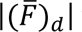 using the cumulative distribution function *F*_(2, 230)_. Lastly, the surge height^27^ representing signal-to-noise ratio of periodic signals is computed by -log10(*P*-value), as shown in the y-axis of the top panels in Fig. 3a,b and Extended Data Fig. 3a,b.

### Gantt charts and sROI mean phases

For each task, a Gantt chart was created by converting the surge profiles of 180 sROIs in each hemisphere into grayscale bars (lower panels, Fig. 3a,b and Extended Data Fig.3a,b, lower panels). Each bar shows the time range where the surge height (amplitude) exceeds 2, resulting from -log10(*P* < 0.01). Portions or whole surges with heights below 2 (*P* > 0.01) are not displayed (white background). Each dot on a bar indicates the mean phase, *θ_sROI_*, obtained by averaging the complex *F*-values of *V* vertices within each sROI using

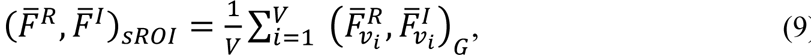

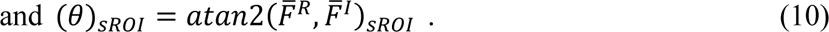

The overall timeline of all Gantt charts is set to a range between 4 and 12 s to encompass all task-related hemodynamic activations.

### Circular statistics

Extended Data Fig. 6 shows the distribution of mean phases across sROIs in each hemisphere for each task. The Watson-Williams test^46,47^ and CircStat (a Matlab toolbox for circular statistics; https://www.mathworks.com/matlabcentral/fileexchange/10676-circular-statistics-toolbox-dir <u>ectional-statistics</u>), was used to assess whether the average of sROI mean phases (*θ_sROI_*) is significantly different between tasks. Given *n*_1_ sROIs for Task 1 and *n*_2_ sROIs for Task 2 (see Supplementary Table 3 for *n*_1_ and *n*_2_):

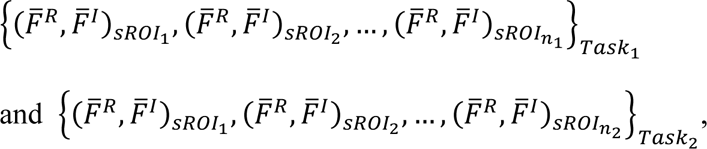

and let *n* = *n*_1_+*n*_2_, the *F*-value of the Watson-Williams test is obtained by:

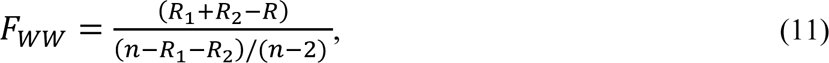

where *R*_1_, *R*_2_, *R* are computed from the radian representations^46^:

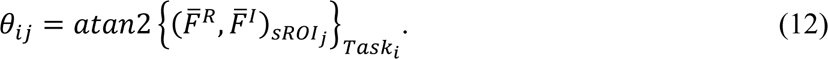

*F_WW_* follows *F*_(1,_ *_n_*_-2)_ distribution approximately.

## Data availability

The data presented in this study can be obtained from the corresponding author upon request.

## Code availability

Custom codes for analyzing phase-encoded fMRI data and traveling waves are included in *csurf* (a FreeSurfer-compatible package) available for download at https://pages.ucsd.edu/~msereno/csurf/ or https://mri.sdsu.edu/sereno/csurf/.

## Acknowledgements

This research was supported by the University of Macau Development Foundation (EXT-UMDF-014-2021), University of Macau (MYRG-CRG2024-00047-ICI, MYRG2022-00265-ICI, MYRG2022-00200-FAH, CPG2023-00016-FAH, CRG2021-00001-ICI, CRG2020-00001-ICI, SRG2019-00189-ICI), Macau Science and Technology Development Fund (FDCT 0001/2019/ASE), and National Institute of Health (R01 MH081990 to M.I.S and R.S.H).

## Author contributions

R.S.H., C.J.G., U.M.L., T.I.L., J.H.A., M.I.S., D.L., and V.L.C.L. contributed to the conceptualization of the research and wrote the paper. R.S.H., C.J.G., C.T.L., and C.U.C. performed experiments. R.S.H., J.H.A., and M.I.S. developed the methodology for data analysis and visualization. R.S.H., C.J.G., and J.H.A. analyzed the data.

## Additional information

**Extended data** is available for this paper at [TBD]

## Supplementary information

The online version contains supplementary material available at [TBD]

**Correspondence and requests for materials** should be addressed to Ruey-Song Huang

---

**Extended Data Fig. 1.**
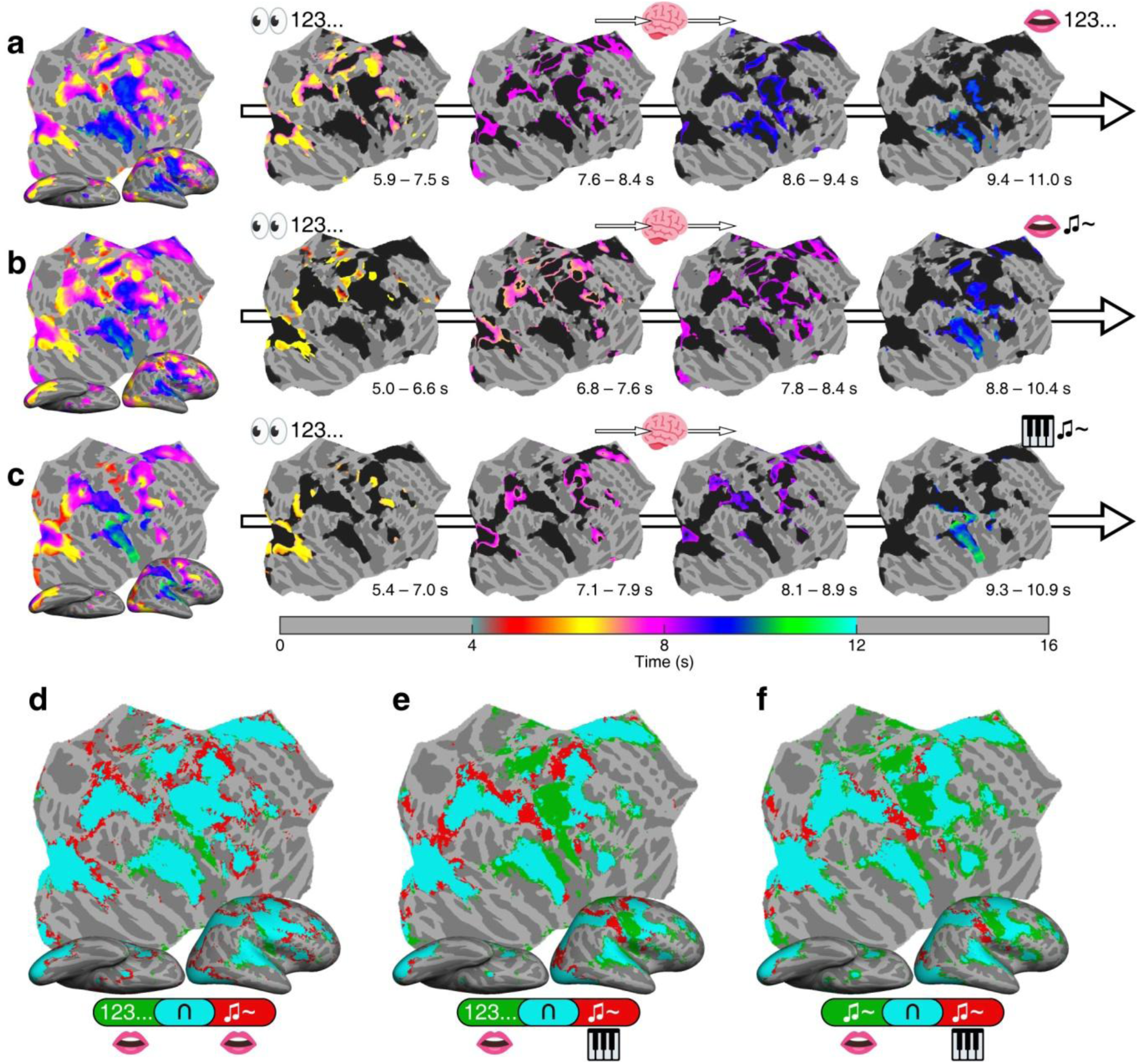
**a**, **b**, and **c**, Group-average maps of phase-encoded activations (*n* = 21, F_(2,40)_ = 5.18, *P* < 0.01, cluster corrected) in the right hemisphere for the digit reading-reciting task, note reading-singing task, and note reading-playing task, respectively. **d**, **e**, and **f**, Conjunction maps. Green or red: regions activated by a single task; Cyan: regions activated by both tasks.

**Extended Data Fig. 2.**
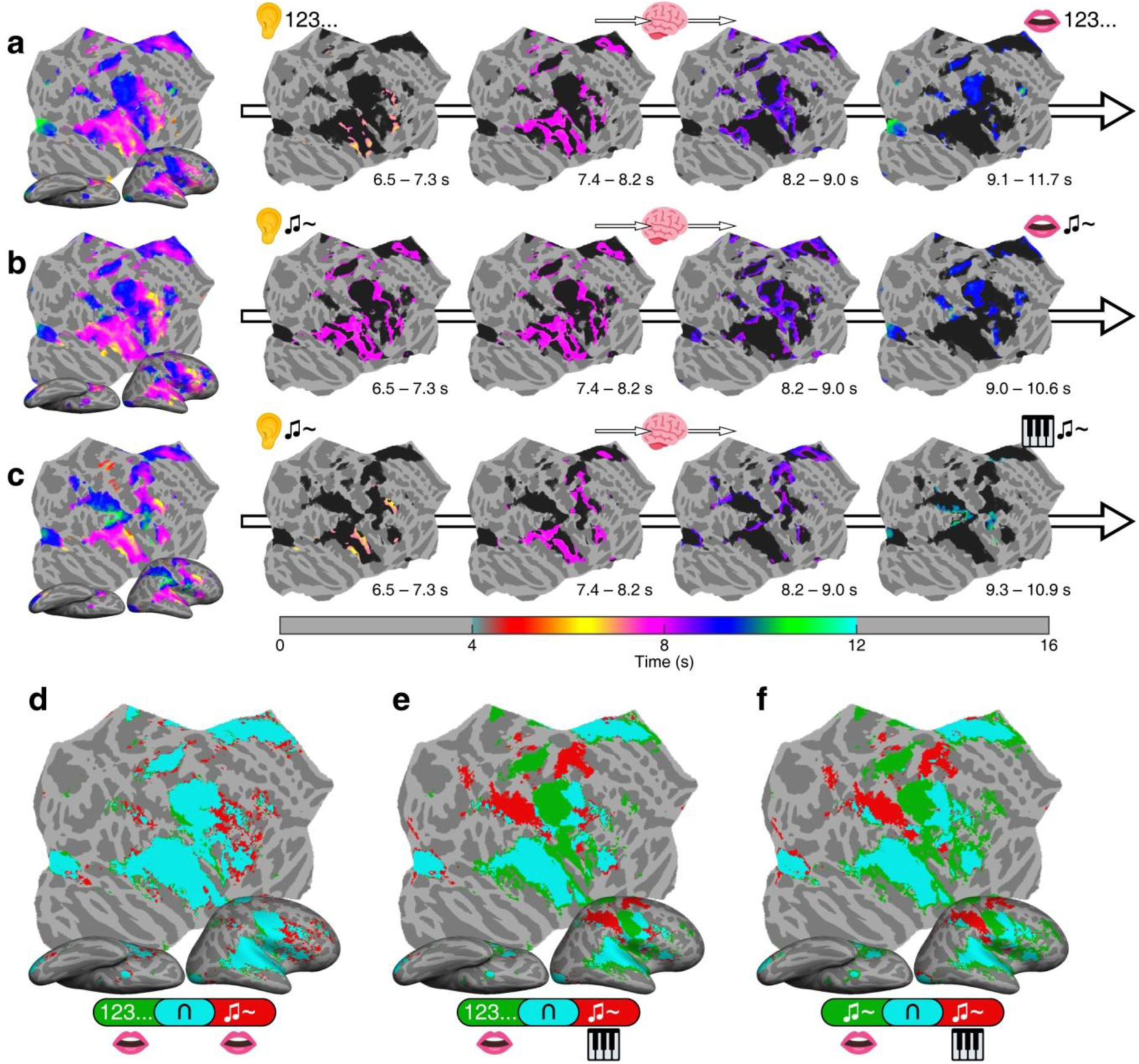
**a**, **b**, and **c**, Group-average maps of phase-encoded activations (*n* = 21, F_(2,40)_ = 5.18, *P* < 0.01, cluster corrected) in the right hemisphere for the digit listening-reciting task, note listening-singing task, and note listening-playing task, respectively. **d**, **e**, and **f**, Conjunction maps. Green or red: regions activated by a single task; Cyan: regions activated by both tasks.

**Extended Data Fig. 3.**
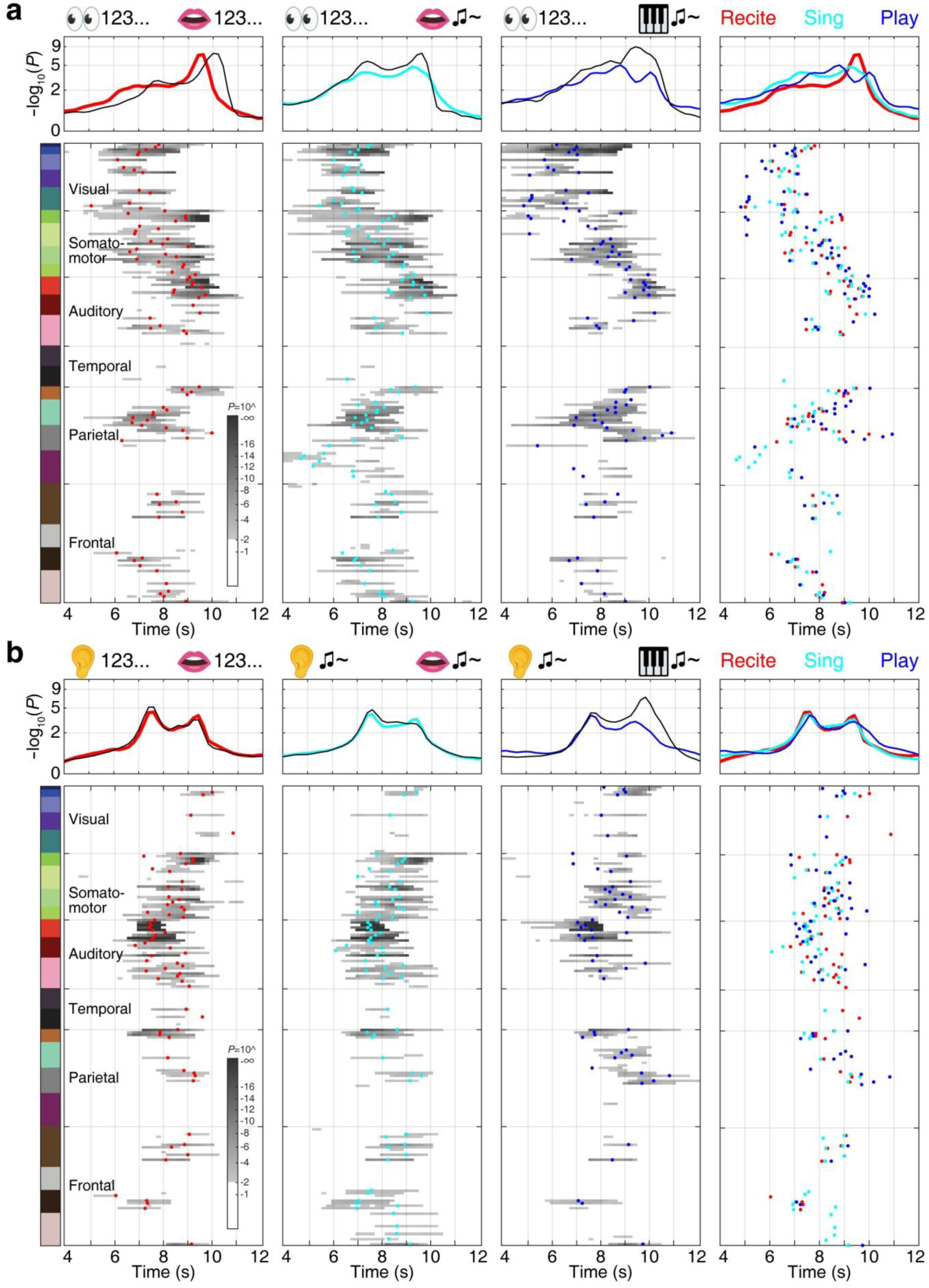
**a**, Surge profiles (upper panels) and Gantt charts (lower panels) of activations in the right hemisphere during the reading-reciting (red), reading-singing (cyan), and reading-playing (blue) tasks. Each black curve represents the surge profile of activations in the left hemisphere for each task. **b,** Surge profiles (upper panels) and Gantt charts (lower panels) of activations in the right hemisphere during the listening-reciting (red), listening-singing (cyan), and listening-playing (blue) tasks. All conventions follow those of Fig. 3a.

**Extended Data Fig. 4.**
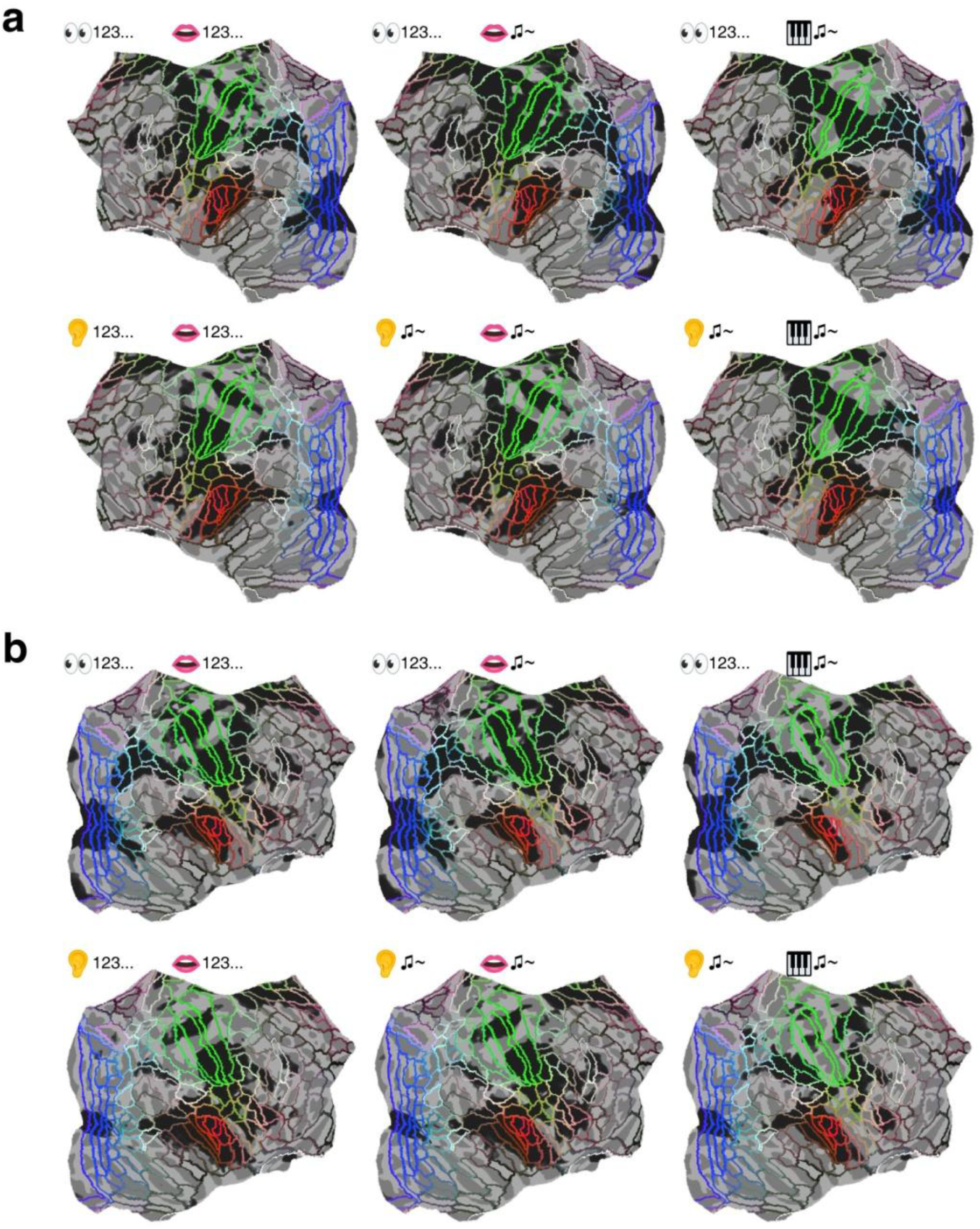
Maps of activations (black regions) overlaid with borders of surface-based regions of interests (sROIs), delineated based on HCP-MMP1.0 parcellation^43^. **a,** Left hemisphere. **b,** Right hemisphere.

**Extended Data Fig. 5.**
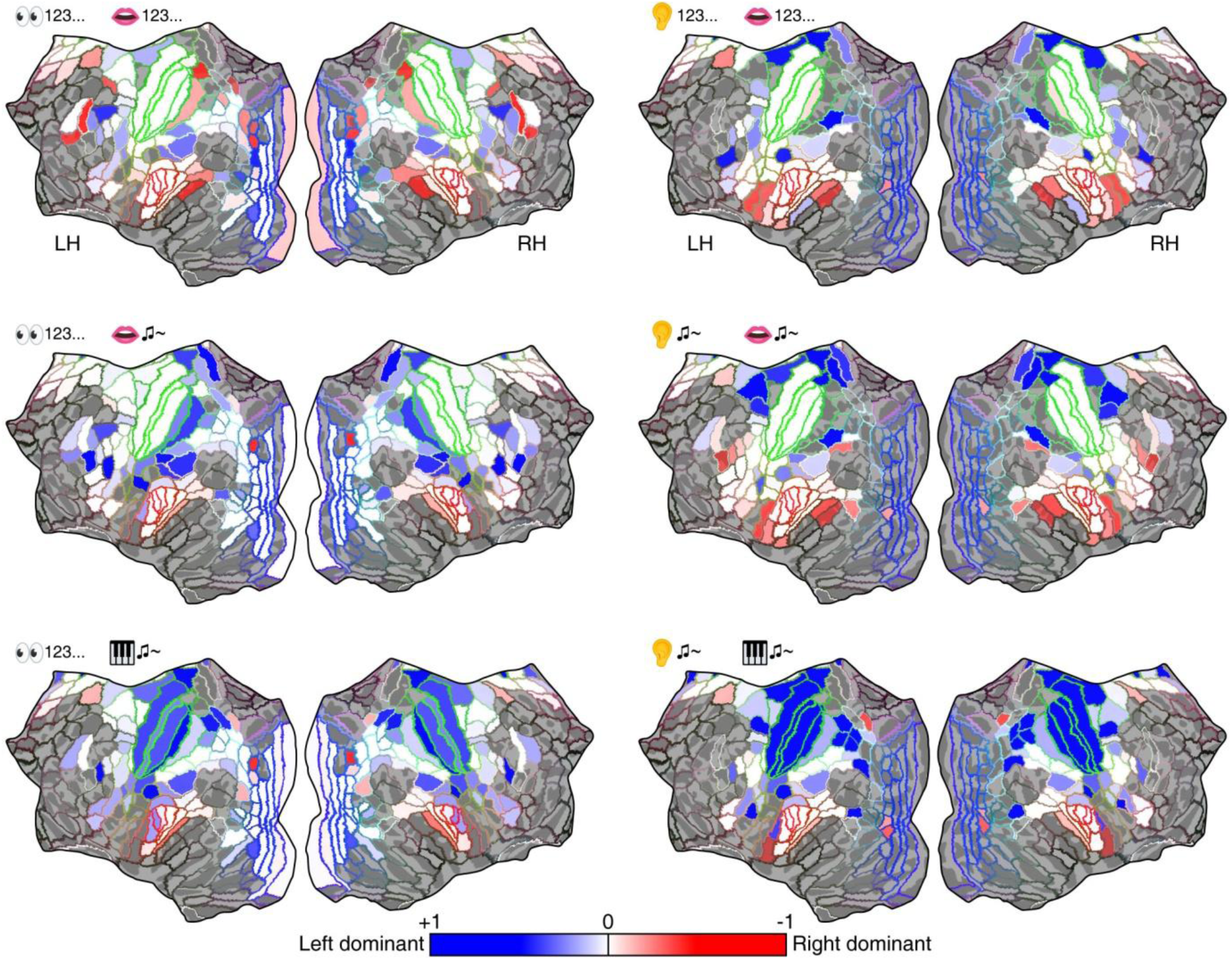
Maps of laterality index (LI) in sROIs delineated in Extended Data Fig. 4. See Supplementary Table 3 for LI values. Corresponding sROIs between hemispheres are rendered with the same color. LH: left hemisphere; RH: right hemisphere.

**Extended Data Fig. 6.**
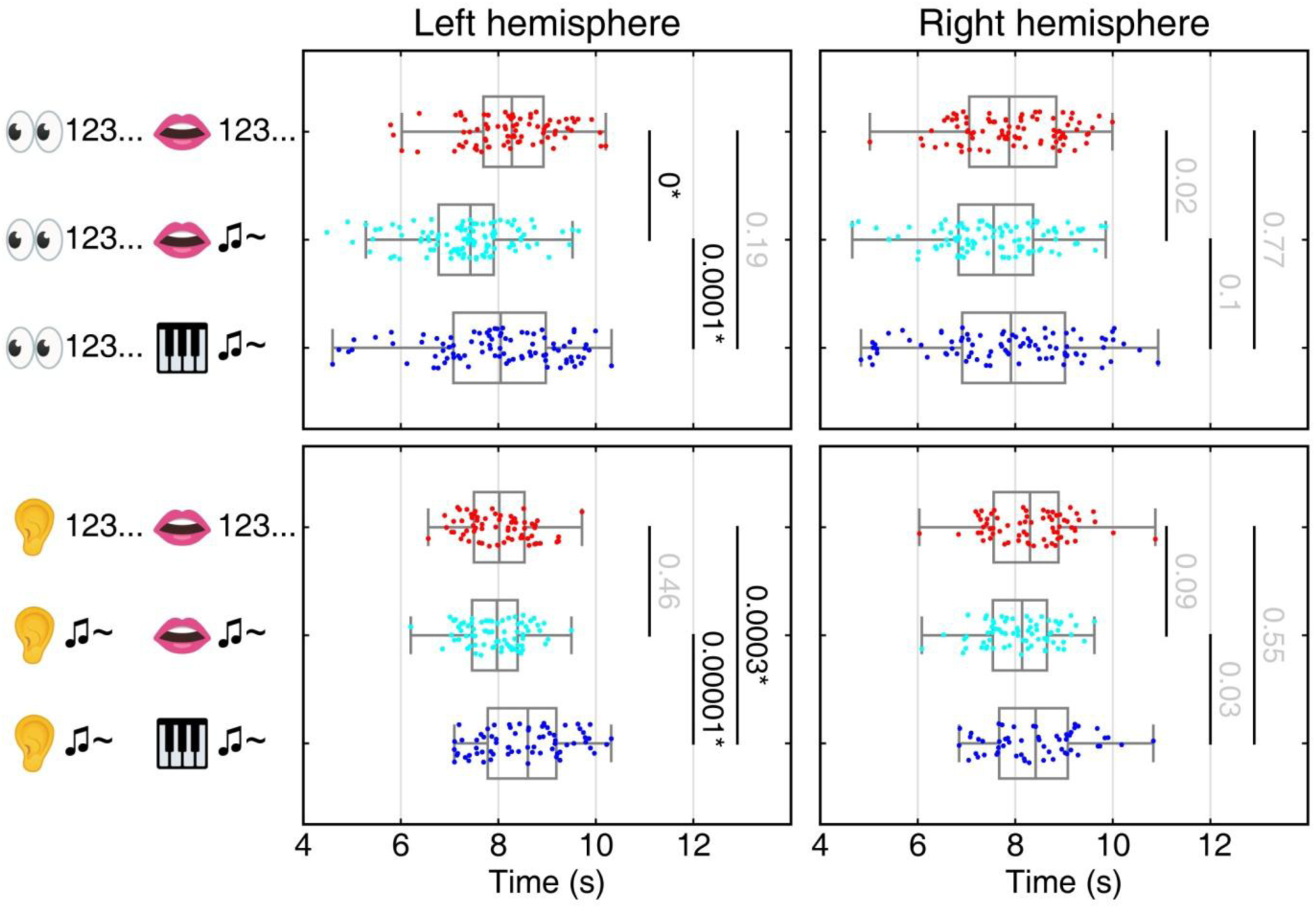
Distributions of sROI mean phases (*θ_sROI_*) in each hemisphere. Black numbers with * indicate the *P*-values for significant difference between tasks (*P* < 0.01, Watson-Williams test; Methods); gray numbers indicate insignificant difference (*P* > 0.01).

**Supplementary Table 1:** Percentages of overlaps between activation maps.

|  | Left Hemisphere |  |  |  | Right Hemisphere |  |  |  |
| --- | --- | --- | --- | --- | --- | --- | --- | --- |
| Reading tasks | Reciting |  | Singing |  | Reciting |  | Singing |  |
|  | 5% | 95% | 65.7% | 34.3% | 12.6% | 87.4% | 73.1% | 26.9% |
|  | Reciting |  | Playing |  | Reciting |  | Playing |  |
|  | 23.5% | 76.5% | 65.8% | 34.2% | 29.6% | 70.4% | 73.1% | 26.9% |
|  | Singing |  | Playing |  | Singing |  | Playing |  |
|  | 29.6% | 70.4% | 87.6% | 12.4% | 36.8% | 63.2% | 71.7% | 28.3% |
| Listening tasks | Reciting |  | Singing |  | Reciting |  | Singing |  |
|  | 10.4% | 89.6% | 81.9% | 18.1% | 12.2% | 87.8% | 79.9% | 20.1% |
|  | Reciting |  | Playing |  | Reciting |  | Playing |  |
|  | 35% | 65% | 56.3% | 43.7% | 49.5% | 50.5% | 60.3% | 39.7% |
|  | Singing |  | Playing |  | Singing |  | Playing |  |
|  | 35.8% | 64.2% | 61% | 39% | 51.1% | 48.9% | 64.1% | 35.9% |

*Note:* The value in each white cell represents the percentage of vertices with significant activations ( $F_{(2,230)} > 7.1$ , $P < 0.001$ ) for a single task. The value in each shaded cell represents the percentage of vertices with significant activations in both tasks, as shown in the conjunction maps.

**Supplementary Table 2:** Group-average task performance (n = 21).

| Task | Response time (s)<br>(mean ±SD) | Response duration (s)<br>(mean ±SD) | Accuracy (%)<br>(mean ±SD) |
| --- | --- | --- | --- |
| Reading-reciting | 1.11 ±0.2 | 2.40 ±0.39 | 99.30 ±0.79 |
| Reading-singing | 1.11 ±0.18 | 2.82 ±0.53 | 88.52 ±15.22 |
| Reading-playing | 0.92 ±0.17 | 3.24 ±0.57 | 98.89 ±1.40 |
| Listen-reciting | 0.58 ±0.11 | 2.44 ±0.22 | 99.81 ±0.36 |
| Listen-singing | 0.62 ±0.16 | 2.61 ±0.34 | 98.95 ±1.70 |
| Listen-playing | 0.7 ±0.16 | 3.112 ±5 | 87.01 ±11.58 |

*Note:* The response time was measured from the onset of a visual cue prompting overt production.

**Supplementary Video 1.**
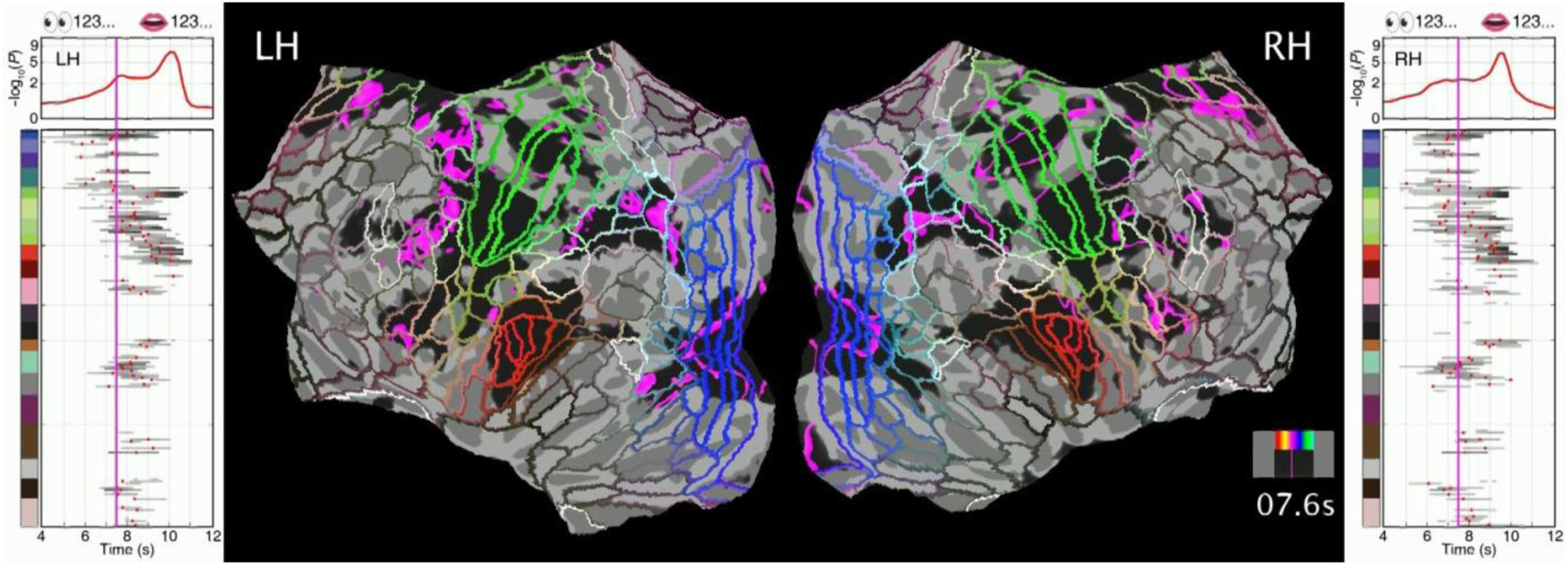
Animated Gantt charts and traveling waves of the digit reading-reciting task.

**Supplementary Video 2.**
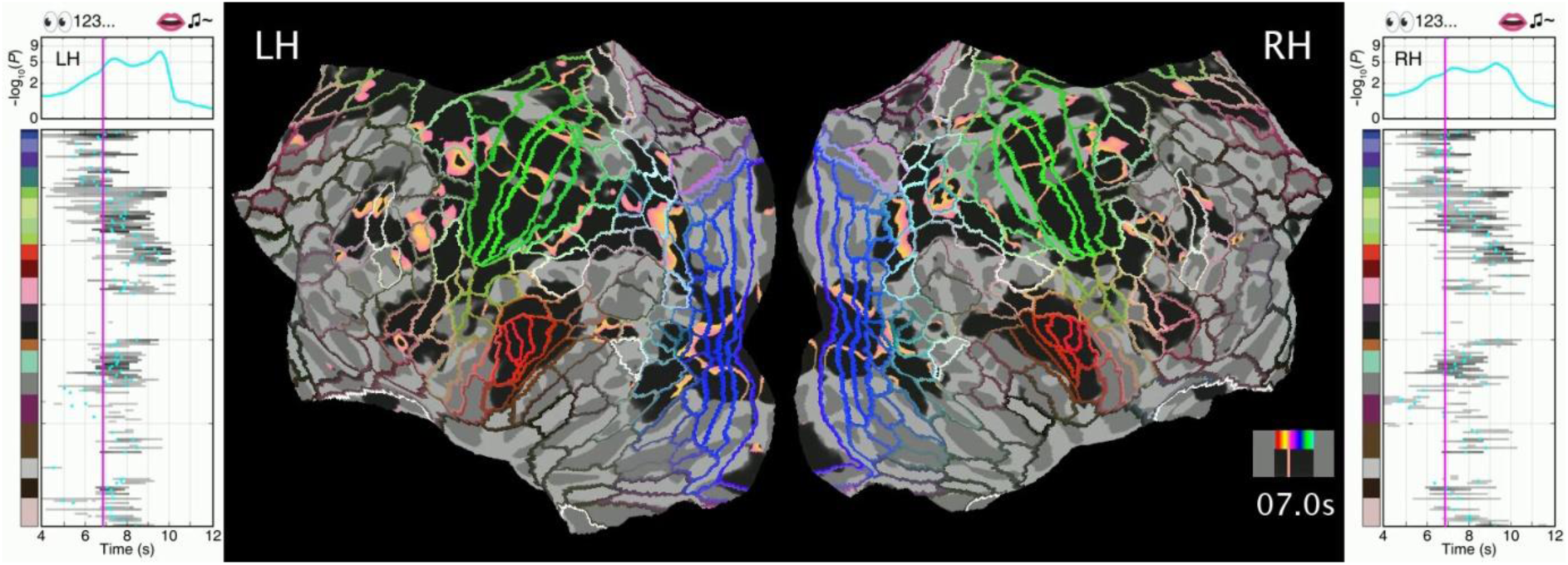
Animated Gantt charts and traveling waves of the note reading-singing task.

**Supplementary Video 3.**
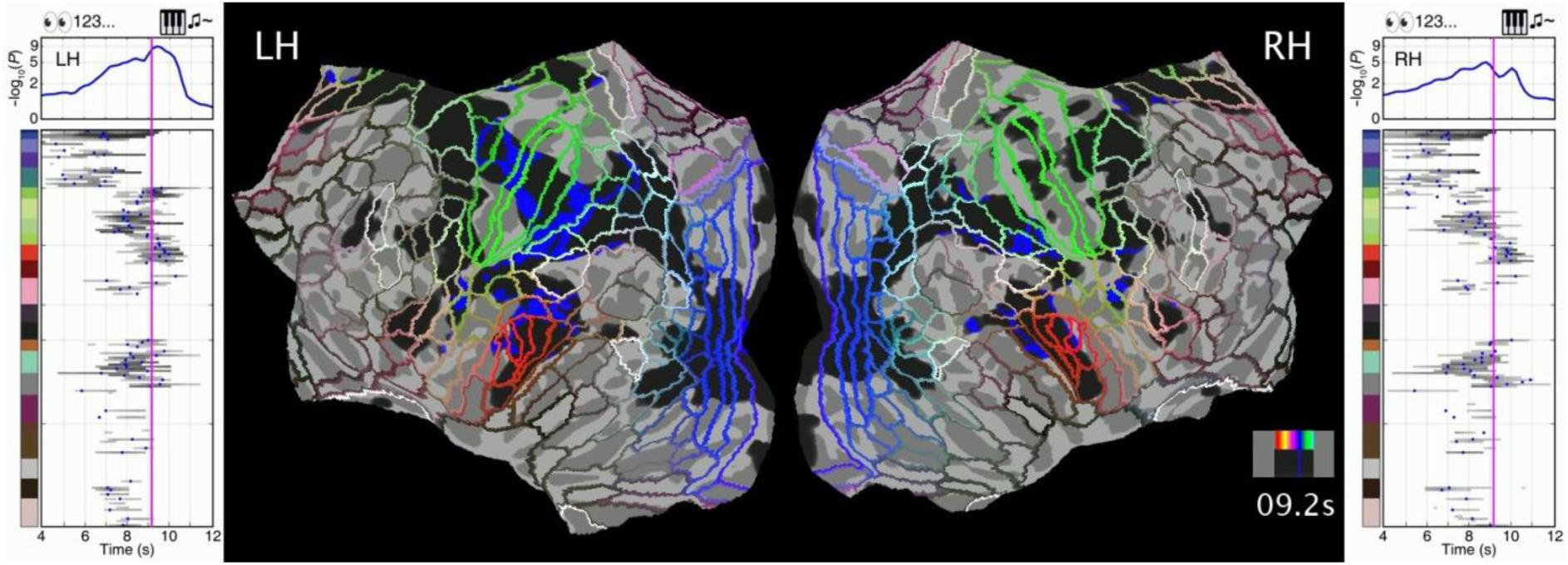
Animated Gantt charts and traveling waves of the note reading-playing task.

**Supplementary Video 4.**
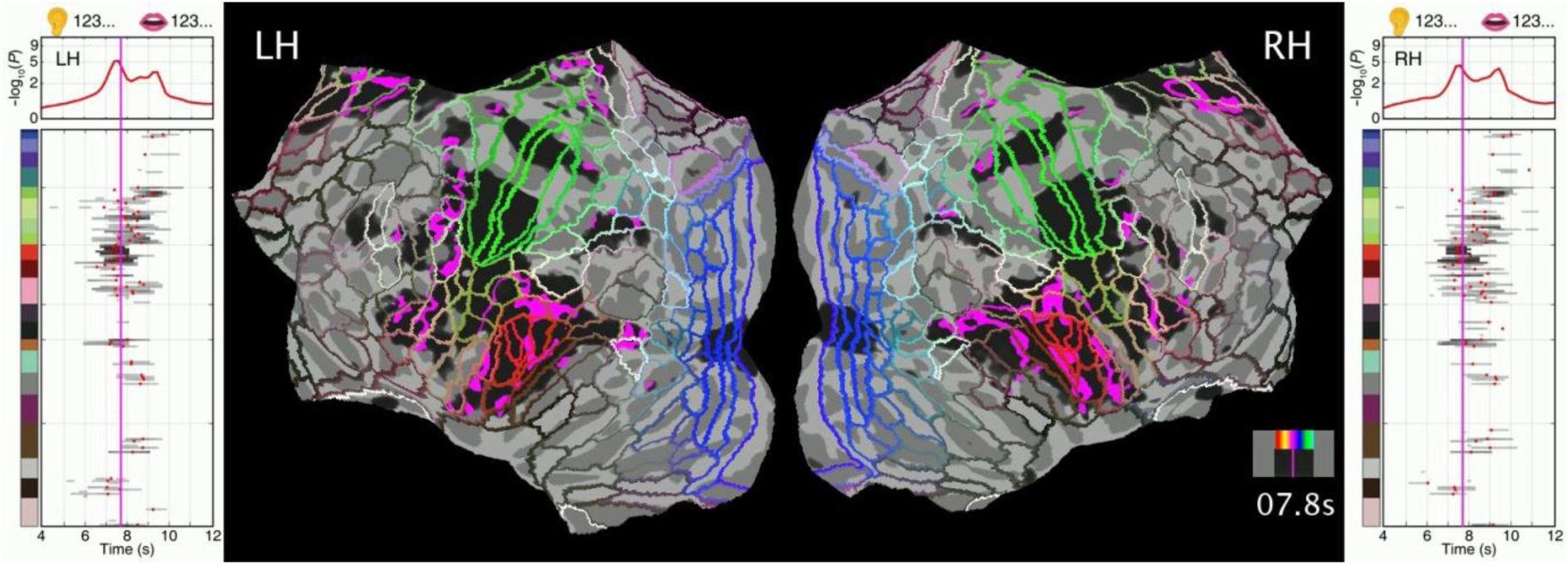
Animated Gantt charts and traveling waves of the digit listening-reciting task.

**Supplementary Video 5.**
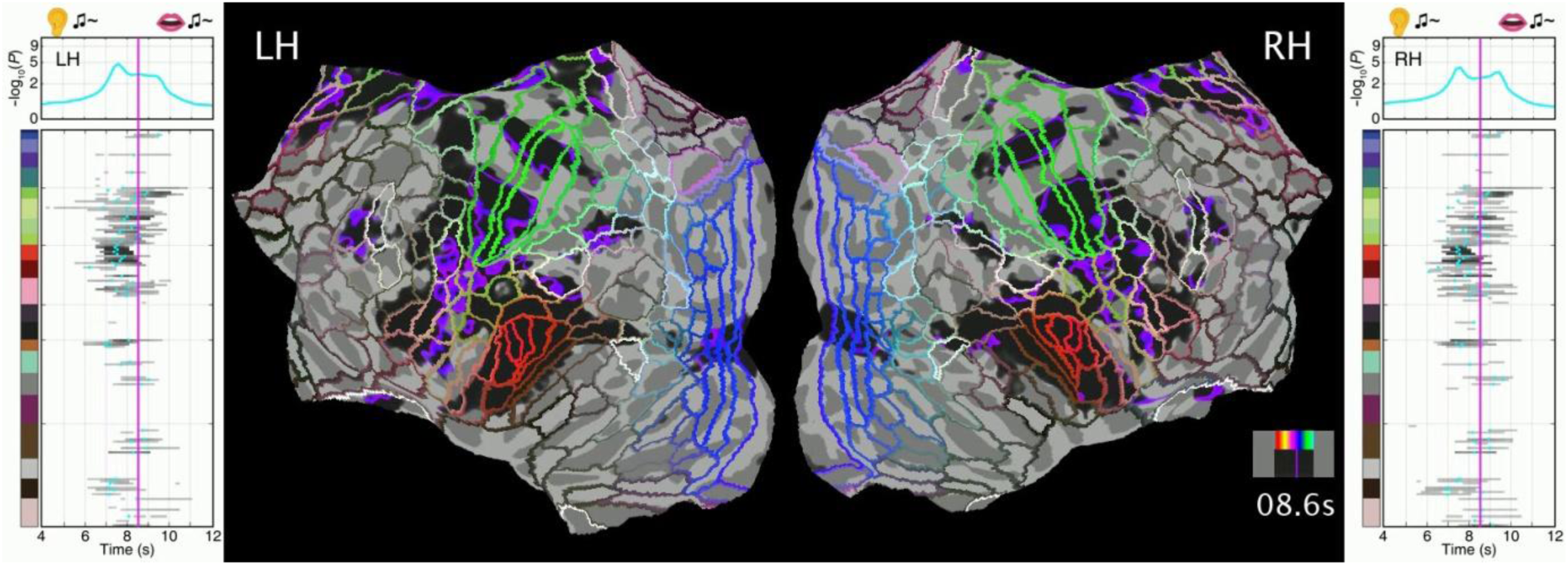
Animated Gantt charts and traveling waves of the note listening-singing task.

**Supplementary Video 6.**
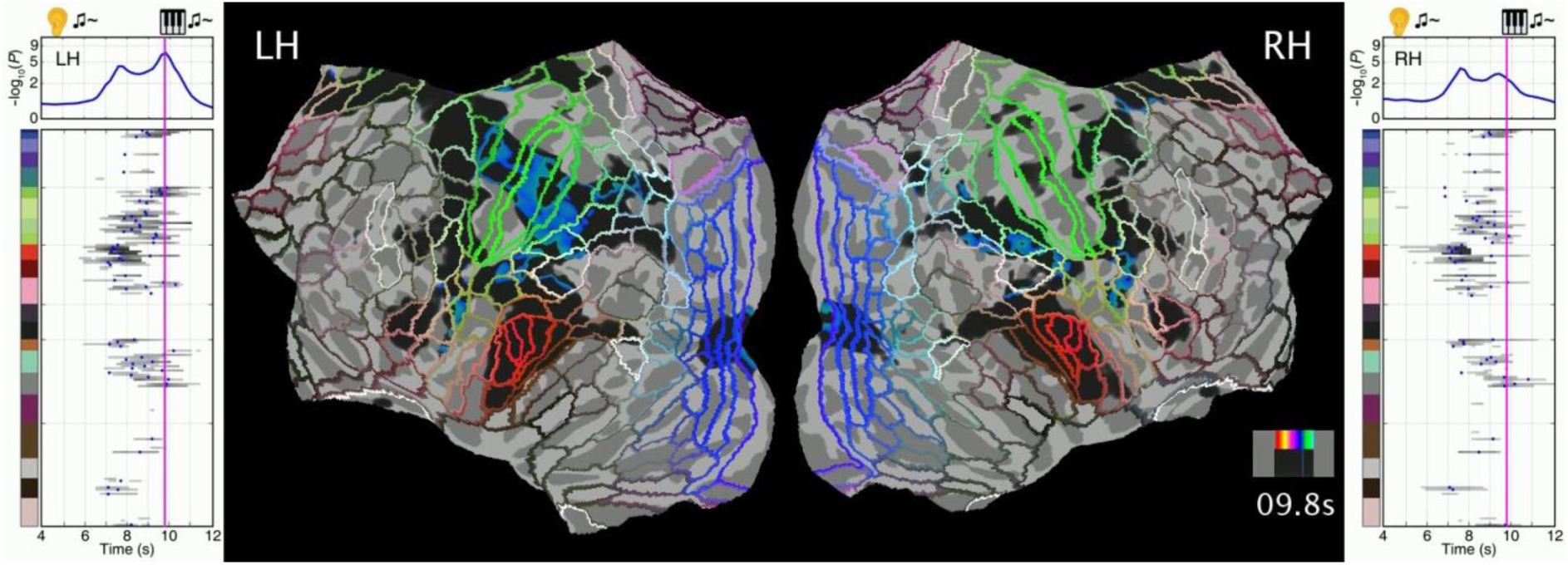
Animated Gantt charts and traveling waves of the note listening-playing task.

